# Identification of transgene insertion sites via short- and long-read whole genome sequencing

**DOI:** 10.64898/2026.02.05.704064

**Authors:** L Kaplan, SJ Edgerton, BD Mahoney, CA Ray, TA Reh

## Abstract

Transgenic mouse lines are essential to uncover organ or system level genotype-phenotype relationships. The generation of such lines via transgene addition may lead to the insertion into unknown genomic loci potentially leading not only to the disruption of native genes but also attenuation of transgene expression. Additionally, this often results in the inability to determine transgene zygosity which in turn complicates breeding and interpretation of experimental results. In this study we present two whole genome sequencing based pipelines that allow the identification and genotyping of even complex multi transgenic inserts. As they use widely available reagents and bioinformatic tools, they can easily be applied to develop genotyping strategies in potentially any species.

## Introduction

Transgenic mouse lines are currently indispensable for modern biomedical research as they enable the study of a variety of gene functions on a complex systemic level. Some of the most common modifications include the knockout or overexpression of genes of interest. Knockouts are usually site-specific, even if a cell type specific conditional recombinase needs to be inserted. While gene insertions, be it fluorescent reporters or ectopic overexpression cassettes, can be specifically targeted to certain loci, classic techniques lead to the insertion in an unknown location. Often, this does not pose a direct problem as long as the insertion occurs in intergenic space with open chromatin. However, in some cases it might lead to unwanted effects like the disruption of native gene structure, multiple insertion sites leading to a dosage effect or even silencing of the transgene [1], [2], [3].

Furthermore, genotyping for homozygous animals is highly complicated without knowledge of the genomic context of the insertion site.

In this study we present two whole genome sequencing based approaches to identify the insertion sites of three separate transgene cassettes of interest and a PCR-based genotyping strategy. The first method is based on short read sequencing and was inspired by the joint profiling of the epigenome and lentiviral integration site analysis (EpiVIA) [4]. Due to the complexity of the mouse line and insert sequence redundancy, we further employed long-read based whole-genome sequencing coupled with a search for structural variations. Both methods utilize standard library preparation and sequencing technologies available in many core facilities as well as free, widely used software.

The mouse line we used to test our protocols contains a tamoxifen-dependent Cre recombinase (CreER) under the control of a glial-specific promoter, that drives the activation of a tetracycline-controlled transactivator (tTA) in the Rosa locus. This in turn activates three separate cassettes: One drives the overexpression of Ascl1 and GFP, one drives expression of Atoh1, and a third contains a combination of Islet1 and Pou4f2 (Fig. 1A). While the locations of the CreER (Glast locus) and tTA (Rosa locus) are known, the rest of the transgenes were generated using pronuclear injections leading to random insertion [5], [6], [7]. Using whole genome sequencing, we were able to identify the genomic loci of each of the transgenes.

**Figure 1:**
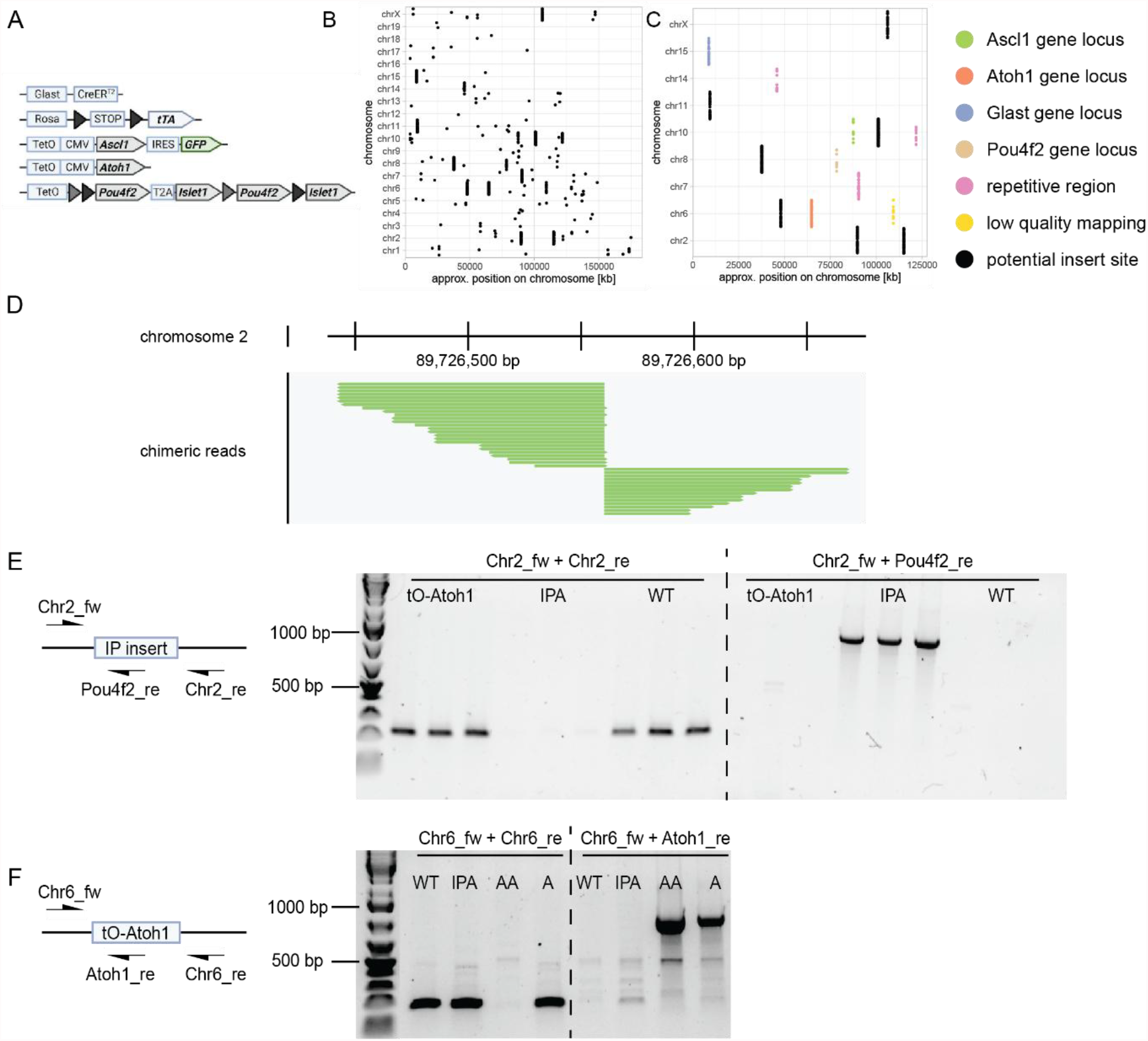
A) Summary of gene inserts of the IPA + Atoh1 mouse line adapted from [9]. B) Chromosomal locations of chimeric reads, that map in part to one insert and one chromosome. C) Chimeric reads as in B but filtered to remove sites with low coverage (< 5 reads) and colored according to known features. D) IGV track of chimeric reads mapping to both sides of the IP insert reveal its insertion site on chromosome 2. E) Primer design and PCR results of the genotyping for the IP insert. Chr2_fw + chr2_re, flanking the suspected insert site, amplify the wildtype sequence in WT and tetO-Atoh1 mouse, but fail to produce the long and complex full-length insert in the IPA mice. Chr2_fw + Pou4f2_re, which binds inside the insert, produces bands of the right size for IPA mice and no bands for WT or tetO-Atoh1. In this case all IPA mice were homozygous for the insert. F) Primer design and PCR results of the genotyping for the tetO-Atoh1 insert. Chr6_fw + chr6_re, annealing on both sides of the putative insert site, yielded wildtype bands for WT, IPA and one tetO-Atoh1 (A) mouse, but not in the tetO-Atoh1/tetO-Ascl1 (AA) mouse. Chr6_fw + Atoh1_re produced bands of the right size only in AA and in tetO-Atoh1 (A) mice. This revealed that this tetO-Atoh1/ tetO-Ascl1 mouse was homozygous for tetO-Atoh1 insert, while the tetO-Atoh1 mouse was heterozygous.

This allowed us to develop a PCR genotyping strategy for transgenes of these unknown insertion loci.

## Results

### Whole genome sequencing with EpiVIA identifies IP and Atoh1 insert loci

To identify as many transgene insertion sites as possible from a single sequencing run, we isolated DNA from the liver of a mouse carrying Glast-CreER, tetO-CMV-Ascl1-GFP (tetO-Ascl1), tetO-CMV-Atoh1 (tetO-Atoh1) and Islet1-Pou4f2 (IP) inserts (Fig. 1A). We performed short read Illumina sequencing to a coverage of >50x of the whole genome. The integration site analysis was inspired by EpiVIA [4] and is based on the identification of chimeric fragments, i.e. reads that map in part to the native chromosome and in part to an insert of interest. In their analysis, there are three meaningful cases in which reads can be informative for integration site analysis: In Case C one end of a paired read fragment maps to the chromosome and one to the insert. Similarly, in Case D, one end maps completely to the chromosome, but the other overlaps the boundary between insert and chromosome. Lastly Case E describes the opposite, where one end overlaps the boundary between insert and native genome, while the other completely aligns with the insert. Because in all these cases one has information of chromosome locus and insert from the same fragment, they reveal the location of the integration. For this type of analysis, the aligner software needs to be able to map any given read to distinct locations on the chromosome and report these as chimeric fragments. For this purpose the Burrows–Wheeler aligner (bwa) [8] is a popular choice. Additionally, it is necessary to create a custom reference genome that includes the sequences of the transgenes of interest. In lentiviral integration site analysis, the sequence is typically known, since it is contained between the LTRs. However, classic transgenic mouse creation via pronuclear microinjection of genetic material can lead to the insertion of not only the desired construct but also part of the plasmid construct of origin or an array of inserts. Here we approximated the insert sequence by reconstructing the respective known gene elements in the correct order for tetO-Ascl1, tetO-Atoh1 and IP and added them to the reference mouse genome as an artificial chromosome. We used bwa to map the sequenced genome to the custom reference. It reported reads of chimeric fragments with the “SA” tag and noted which chromosomes they map to, allowing us to pull out all read pairs that map in part to the chromosome and in part to an insert. This left us with around 10 000 such reads mapping to 125 genomic loci (Fig. 1B). Removing regions that were covered by 5 or fewer reads, eliminated the background and retained 19 well defined genomic regions of interest. Closer inspection of these revealed that some were located at the sites of the native genes that were also present in our transgene cassettes. This occurred if, for example, a fragment mapped to the native Ascl1 locus as well as to the insert. In this case the fragment was tagged as chimeric, although it probably stemmed from a pure insert fragment, that also contains Ascl1 sequence. We found such sites for genomic loci of Glast (Slc1a3), Ascl1, Atoh1, Islet1 and Pou4f2 (Fig. 1C). Further regions could be filtered out due to low quality mapping or repetitive sequences leaving 11 potential insertion sites. Looking at the coverage of the region, we determined that a site on chromosome 2 (mm10: chr2 89726484) was highly likely to harbor the IP insert (Fig. 1D). The chimeric fragments here cover both ends of the insert with reads mapping to tetO on one side and to FRT on the other end. Since the FRT element is unique to the IP insert, we were confident to have identified the first insertion site. We designed primers that bind up and downstream of the putative insertion site and a third primer that specifically binds in the Pou4f2 gene inside the cassette. PCR results from animals with known genotype (earlier zygosity-agnostic genotyping) with the surrounding primers yielded the expected short band (~200 bp) for wildtype animals and no band for mutants (Fig. 1E). Although a longer mutant band (>6 kb) would be expected, the PCR conditions we used did not allow its amplification. To allow the positive identification of the mutant allele, we combined the chromosome 2 forward primer with Pou4f2 reverse, which in turn yielded a clear band (~1500 bp) for IP + tetO-Ascl1 (IPA) animals, but not for wildtype or tetO-Ascl1 mice. A band in both reactions is expected for heterozygous animals, while a band in only one or the other reaction represents homozygous animals. Thus, the future genotyping strategy will entail the identification of wildtype alleles with the surrounding primers and combine it with the primer binding to Pou4f2 to test for mutant alleles.

Unfortunately, there were no chimeric fragments uniquely identifying either the tetO-Ascl1 or tetO-Atoh1 sites. This was because both sequences carry identical tetO-CMV sequences in the 5’ region and similar poly adenylation signals at their respective 3’ ends. Subsequently, we systematically investigated all 10 remaining potential regions designing primers as previously described for the IP site and we were successful in finding the tetO-Atoh1 site on chromosome 6 (mm10: chr6 47968717) (Fig. 1F)

### Long-read WGS and structural variation analysis identifies tetO-Ascl1 insert locus

All attempts to identify the tetO-Ascl1 site, however, failed, due to the ambiguity of the detected chimeric fragments. To overcome this roadblock, we isolated genomic DNA from a mouse carrying only tetO-Ascl1 (Fig. 2A) and used Oxford Nanopore Technology based long-read whole genome sequencing. This eliminated our reliance on chimeric reads, as we can expect individual long reads covering the whole insert plus the flanking chromosomal sequences without any ambiguity.

**Figure 2:**
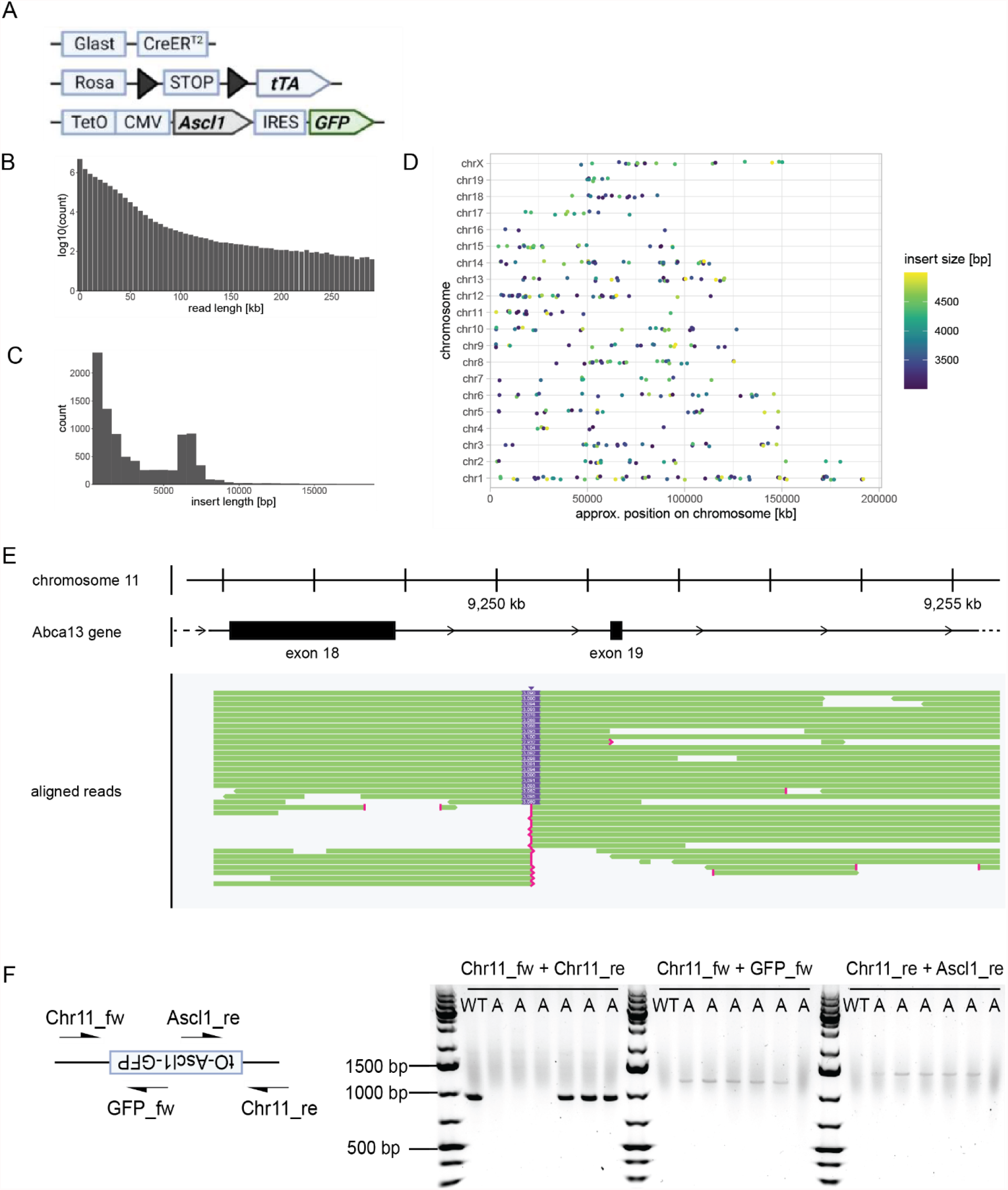
A) Summary of gene inserts of the tetO-Ascl1 mouse. B) Distribution of read length achieved by long-read whole genome sequencing. C) Distribution of insertion lengths detected by calling structural variations between the sequenced mouse and the reference genome mm39. D) Chromosomal locations of all detected insertions between 3 and 5 kb color coded by length. E) Alignment track of the genomic locus on chromosome 11 showing reads covering the roughly 3.1kb sized insertion (purple) that hosts the tetO-Ascl1 transgene, which is located inside an intron of the Abca13 gene. F) Primers were designed to amplify the whole locus or bind specifically to known parts of the insert. Insert flanking primers produced the correct wildtype band in Bl6 animals (WT) and in 3 animals from the tetO-Ascl1 line (A). If flanking primers were combined with either GFP or Ascl1 targeting primers, they only yielded bands for animals from tetO-Ascl1 litters. In this example the first 3 individuals from the tetO-Ascl1(A) line are homozygous for the insert, since no wildtype band is present. The following two animals are heterozygous and produce the wildtype as well as GFP, and Ascl1 bands. The last animal is negative for the tetO-Ascl1 insert.

Indeed, whole genome sequencing on a PromethION produced millions of reads that ranged from few to hundreds of kb in length (Fig 2B). We hypothesized that by comparing our transgene mouse line with a wildtype, any big insert would be treated as a structural variant the same way if one would compare different individuals or strains. Thus, we used the long-read aligner minimap2 [10], [11] to map reads to the reference genome without the addition of custom sequences. Subsequently, we used kled [12] to call structural variants that contain amongst others insertions, deletions, indels or inversions. The resulting variant call file (vcf) reports the chromosomal position of any structural variant as well as its sequence. As the sequenced mouse and the wildtype are from different strains, we expected to find a large number of small variations especially in intergenic regions. Indeed, we found a total of 87566 (Fig. 2C) variants, however even after filtering only for insertions of the desired size (3000-5000 bp) we were surprised to see that 946 (Fig. 2D) variants remained. This high number prohibits checking whether each called insert contains transgene sequence manually. Furthermore, a BLAST search that would reveal sequence identity for multiple queries is limited to a set amount of basepairs that in our case was far exceeded. However, using the BLAST Command Line Applications [13] does not pose such limit. Using the insert sequences called by kled as the query against the refseq transcriptome database, we received exactly one hit that matched the Ascl1 sequence and was located on chromosome 11 (mm39: chr11 9250364) (Fig. 2E). Looking at the sequence of the insert, we confirmed the presence of all elements of the transgene. Tracing it back to the region, it was covered by reads with lengths up to 30 kb, we looked at one representative read with a length of 5.2 kb that revealed the complete context of our insert. The cassette was inserted on the antisense strand between two exons of the Abca13 gene. Interestingly, this region was indicated in our previous short-read sequencing, but was, however, deprioritized in our efforts due to some ambiguity and low read coverage as described above.

Again, we designed flanking primers to cover the whole region and one primer that binds specifically to either Ascl1 or GFP. However, even after various attempts to optimize the PCR reaction in terms of primers, temperatures, additives and polymerases, we were not able to specifically amplify either the wildtype or mutant region. Upon further investigation, we found that the 5’ end of this chromosomal locus not only contains long stretches of repeats but is also rich in AT bases. This low complexity was further aggravated by the repetitive nature of the tetO element at the genomic 3’ end. Taken together, this not only complicated the design of specific primers but also the reliable amplification of a specific product, resulting in various bands of different sizes for both genotypes. Subsequently, we tested polymerases that were designed to robustly amplify difficult or very long targets and were able to get the desired band using RepliQa HiFi ToughMix (Fig. 2F). However, while it was able to reproducibly amplify the wildtype locus, we were still not able to produce the full length insert from flanking primers. Finally, we combined the primers that bind chromosome 11 with either Ascl1 or GFP targeting primers, respectively and were thus able to confirm the location, orientation and identity of the tetO-Ascl1 insert (Fig. 2F).

## Discussion

Many transgenic mouse lines that carry gene insertions were generated using pronuclear injection of genetic material and lead to a random insertion into the genome. Not only can this cause the disruption of native genes, it can also lead to suboptimal transgene expression. Additionally, while the detection of the transgene is possible with primers that bind to its known sequence, this method can not determine the zygosity. This in turn can lead to a difficult interpretation of experimental results if, for example the introduced mutation is recessive or otherwise dependent on gene dosage. Some methods have been developed to discern animals with two mutant alleles from those carrying only one. One such protocol relies on the assumption that the gene expression in animals with two alleles should be significantly higher and thus should be measurable by quantitative PCR. However, this requires the possibility to design specific and reliable primers and a discernable difference in expression between the alleles [14], [15]. Another study suggested using unique restriction sites that are introduced on the transgene in addition to the native sequence that is to be overexpressed [15]. An amplification with primers that bind to this native sequence, would amplify both wildtype and mutant site, but only the mutant site would be digested. By comparing digested and undigested DNA products, one can deduce the zygosity. While this method is cost effective and reliable, it is only feasible if specific restriction sites are present and the exact insert sequence is known. Furthermore, neither of the previous methods reveal the genome context of the insert locus, thus precluding any insights in terms of the influence on native gene structure. In contrast, targeted sequencing by proximity ligation, which is inspired by genome architecture methods like HiC, can deliver this information in an unbiased approach [16]. Briefly, DNA is crosslinked, digested and ends ligated leading to circular, closed DNA loops, some of which contain the transgene as well as parts of chromosomes that were in close physical proximity during crosslinking. Using primers that bind in the insert sequence and amplify the whole DNA loop one can get a library that, after sequencing, reveals in which general region of the genome the insert was located. This method allows not only the identification of the haplotype but also of insertion locus. However, it requires specialized DNA and library preparation as well as a sophisticated bioinformatic pipeline.

In this study we presented two strategies based on whole genome sequencing to identify insert locations even in a complex multitransgenic mouse line. The short-read sequencing method that was inspired by EpiVIA requires relatively precise knowledge of the insert sequence as it relies on the identification of chimeric fragments that need to fall exactly at the border between chromosome and insert. Library preparation and bioinformatic pipeline use standard procedures with minimal basic text processing and should thus be accessible to most labs. However, this method requires some manual exploration of read sequences and fails with lines that carry multiple inserts especially if they exhibit sequence redundancies.

These drawbacks are mostly alleviated when using long-read sequencing, which produces reads of tens or even hundreds of kilobases in length and as such capture even the longest of inserts in addition to the chromosomal context. Because the whole cassette sequence is captured, there is no ambiguity even for multiple transgenes with highly similar building blocks. Additionally, only a minimal knowledge of sequence is necessary. While for EpiVIA, it is essential to know the cassette elements at the border of the insert, the variant call method only requires information about any unique feature. The analysis involves slightly more experience in bioinformatic analysis, as it relies on several command line tools, but it has the advantage of being less (wet lab) labor intensive without the need to check individual regions or reads.

Our specific case also highlights the necessity to not only optimize primer design and PCR reaction conditions, but also considering alternative DNA polymerases to achieve a successful genotyping strategy.

Both methods presented in this work are viable options to reliably detect transgenes in mutant model organisms, theoretically in any species, as long as a good reference genome is available. Although library preparation and sequencing might present a major cost, it might be offset by lower animal facility costs due to more efficient breeding and clearer experimental results.

## Materials and Methods

### Data analysis and availability

Bioinformatic analysis was performed on a desktop PC with 16 cores/32 threads with 96 GB of RAM running Ubuntu 22.04. Wherever the software allowed we enabled multithreading to use 30 threads. All plots were generated using R programming language (version 4.4.1). Sequencing data is available at SRA with the accession numbers SAMN55077637 and SAMN55077356. All scripts used for the bioinformatic analysis are available at https://github.com/rehlab/WGS-insert-site.

### Short read library preparation and sequencing

DNA was extracted from the livers of two previously genotyped *Glast-CreER:LNL-tTA:tetO-mAtoh1:tetO-P&I:tetO-mAscl1-IRES-GFP* mice. Genomic DNA was extracted and purified using the DNeasy Blood & Tissue Kit (Qiagen 699506). The quality and quantity of extracted DNA was evaluated using a Nanodrop 2000 spectrophotometer (Thermo Fisher Scientific, Waltham, MA, USA). Four micrograms of purified DNA was sent to Azenta for Short-Read Non-Human Whole Genome Sequencing, including quality control and library preparation with Illumina MiSeq. Sequencing yielded 1 065 million paired end reads with a read length of 150 bp resulting in an average genome coverage of 56.53x. Raw fastq files were preprocessed with fastp [17] with default settings, trimming adapters and performing other QC filtering.

### Bioinformatics pipeline for short read sequencing

To generate a custom reference, we first created a fasta file containing the reconstructed sequences of our transgene cassettes. Subsequently, we generated a genome annotation file (gtf) defining the locations of all elements of our cassettes relative to the fasta file. We then appended both new files to the respective fasta and gtf of the mm10 reference genome. The combined files were used as input to generate the bwa index with the *bwa index* command. The *bwa mem* command was used to align the preprocessed fastq files to the custom reference genome and the output was converted to bam format with samtools [18]. To sort out chimeric reads, we used the output of *samtools view* and piped it into awk for a line by line regex search. Specifically, we looked for SA tags that were followed by “insert” thus yielding chimeric reads that in part cover our artificial chromosomes representing any transgene insert.

The resulting sam file was the basis of the read analysis using an R script. In short, it was treated as a data frame and prepared so that all tags are accessible as columns. For each read we defined cases as suggested by Wang et al. [4] based on the information stored in the “RNAME” (chromosome the read is mapped to), “RNEXT” (chromosome the mate read is mapped to) and “SA” (supplementary alignment) tags [19]. To get a sense of read coverage of any given locus, we defined all reads that fall into a 1000 bp window to be of the same region. Reads were filtered to include regions with more than 5 reads, being of cases c, d or e and not in the location of native genes, repetitive regions or low-quality mapping. The remaining regions were subsequently investigated by PCR. To get chromosomal sequences of regions of interest we used the “Get DNA in Window” tool from the UCSC Genome Browser.

### Long read library prep and sequencing

25 mg of liver was dissected from a mouse with known tetO-Ascl1 genotype and submitted to the Nanopore Sequencing Core of the University of Washington. There, DNA was isolated using the Qiagen Fast DNA Tissue Kit (Qiagen), library prepared (SQK-LSK114-XL, Oxford Nanopore Technologies) and sequenced on a PromethION (FLO-PRO114M, Oxford Nanopore Technologies). The versions of the used tools were: MinKNOW 25.03.7, Bream 8.4.4, Configuration 6.4.10, Dorado 7.8.3, MinKNOW Core 6.4.8. Total sequencing yield was 8.08 Gb (28.9x coverage). We received basecalled, unmapped bam files.

### Bioinformatics pipeline for long-read sequencing

Bam files of reloads were combined using *samtools merge* and converted to fastq format with *samtools fastq* before aligning them to the mm39 reference genome using the longread mode of minimap2 [10], [11] (flags: *-ax map-ont*). The aligned sam file was converted to bam format and used as an input for the variant caller kled [12] resulting in a vcf. Next, we used an R script to filter the variants for inserts between the length of 1000 and 10000 bp to account for incomplete as well as arrayed transgene insertions and exported their sequences as a fasta file. To identify inserts that carry the Ascl1 sequence, we utilized the blast command line suite [13] and started by downloading the cDNA database with update_blastdb.pl with the flags -*-dbtype nucl --decompress refseq_rna*. The inserts fasta file was queried against the database with blastn (flags *-outfmt 6 -evalue 1e-5)*. The result was read into R and filtered for rows containing the GenBank accession numbers of human or mouse Ascl1 (NM_004316.4, NM_008553.5) retrieving one hit located in chromosome 11.

### Animals

All animals were treated and housed in accordance with University of Washington Institutional Animal Care and Use Committee (IACUC) approved protocols. Animals were euthanized by carbon dioxide overdose, followed by cervical dislocation and tissue collection per approved IACUC protocol. Tissue was collected from adult mice over 30 days old. The *Glast-CreER:LNL-tTA:tetO-mAscl1-IRES-GFP, Glast-CreER:LNL-tTA:tetO-mAtoh1:tetO-P&I:tetO-mAscl1-IRES-GFP* and *Rlbp1-CreER:LNL-tTA:tetO-mAscl1-ires-GFP* mice are from mixed backgrounds of C57BL/6 and B6SJF1. The *Glast-CreER, LNLtTA*, and *rtTa* mice are from the Jackson Laboratory. The *tetO-mAscl1-GFP* mice were a gift from M. Nakafuku (University of Cincinnati), the *tetO-Atoh1* mice were a gift from P. Chen (Emory University), the *tetO-P&I* mice were a gift from Professor X. Mu (University of Buffalo) and the *Rlbp1-CreER* mice were a gift from E. Levine (Vanderbilt University).

### Standard PCR

For the identification of the IP and Atoh1 sites, DNA was extracted from ear tags using QuickExtract (Lucigen, QE09050) and 1 µl was used for a PCR reaction using DreamTaq PCR Master Mix (Thermo Scientific, K0182) according to manufacturers’ protocols.

### PCR for complex target

DNA was extracted from the liver and/or ear clippings from the following mouse lines: Bl6 (WT), *Rlbp1-CreER:LNL-tTA:tetO-mAscl1-IRES-GFP*. Genomic DNA was extracted and purified with the Monarch Spin gDNA extraction kit (NEB #T3010S, New England Biosciences). The quality and quantity of extracted DNA was evaluated using a Nanodrop 2000 spectrophotometer (Thermo Fisher Scientific, Waltham, MA, USA). PCR was performed in a total volume of 25 μL, including 12.5 μL of RepliQa HiFi ToughMix (Quantabio, Beverly, MA), 0.375 μL of 10 μM forward primer, 0.375 μL of 10 μM reverse primer, 1 μL of DNA template (concentration >200 ng/μL), and 10.75 μL of nuclease-free water. Thermal cycling included 2 min at 98 °C, followed by 30 (WT Band) or 34 cycles (Mutant bands) of 10 s denaturation at 98 °C, 5 s annealing, 6 s extension at 68 °C, then a final extension step at 68 ° C for 1 min. Annealing temperatures were 63 °C for WT (Chr11_fw+Chr11_re) and GFP mutant band (Chr11_fw+GFP_fw) or 66 °C for Ascl1 mutant band (Chr11_re+Ascl1_re). Amplified products were analyzed using gel electrophoresis on a 1% agarose gel.

**Table 1:**
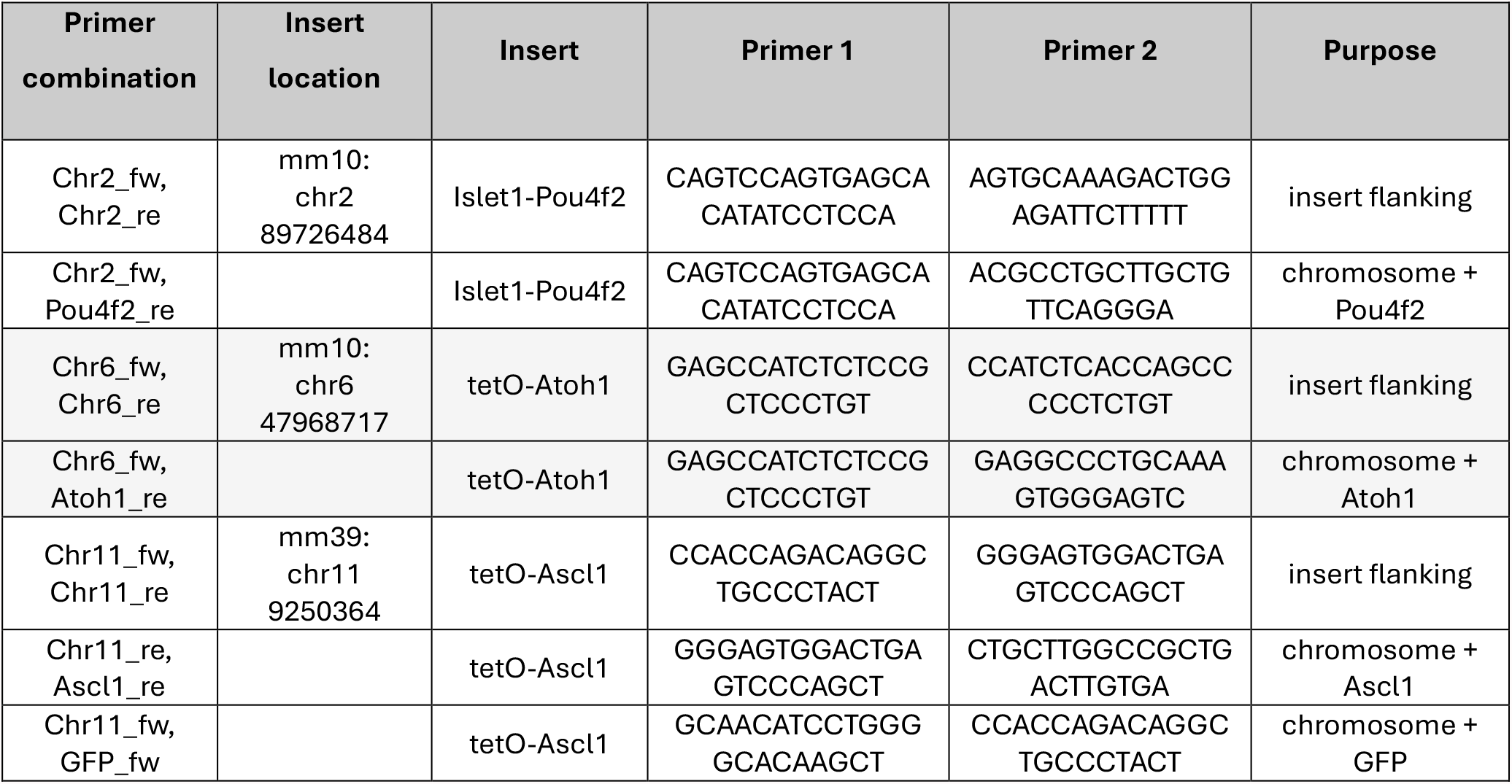
Insert loci and genotyping primers.

## Acknowledgments

We thank the whole team of the Reh lab for great discussions and the Miller lab of the Nanopore Core for the long-read sequencing.

## Author contributions

LK conceptualized the study, designed experiments and bioinformatics pipelines, analyzed data, and wrote manuscript. SJE and BDM performed and analyzed genotyping PCR and edited the manuscript. CAR isolated DNA and coordinated sequencing, edited the manuscript. TAR supervised the project and edited the manuscript.

